# Statistical image processing quantifies the changes in cytoplasmic texture associated with aging in *Caenorhabditis elegans* oocytes

**DOI:** 10.1101/2020.07.30.228270

**Authors:** Momoko Imakubo, Jun Takayama, Hatsumi Okada, Shuichi Onami

## Abstract

**Background:** Oocyte quality decreases with aging, thereby increasing errors in fertilization, chromosome segregation, and embryonic cleavage. Oocyte appearance also changes with aging, suggesting a functional relationship between oocyte quality and appearance. However, no methods are available to objectively quantify age-associated changes in oocyte appearance.

**Results:** We show that statistical image processing of Nomarski differential interference contrast microscopy images can be used to quantify age-associated changes in *Caenorhabditis elegans* oocyte appearance. Max-min Value (mean difference between the maximum and minimum intensities within each moving window) quantitatively characterized the difference in oocyte cytoplasmic texture between 1- and 3-day-old adults (Day 1 and Day 3 oocytes, respectively). With an appropriate parameter set, the gray level co-occurrence matrix (GLCM)-based texture feature *Correlation* (COR) more sensitively characterized this difference than the Max-min Value. Manipulating the smoothness of and/or adding irregular structures to the cytoplasmic texture of Day 1 oocyte images reproduced the difference in Max-min Value but not in COR between Day 1 and Day 3 oocytes. Increasing the size of granules in synthetic images recapitulated the age-associated changes in COR. Manual measurements validated that the cytoplasmic granules in oocytes become larger with aging.

**Conclusions:** The Max-min Value and COR objectively quantify age-related changes in *C. elegans* oocyte in Nomarski DIC microscopy images. Our methods provide new opportunities for understanding the mechanism underlying oocyte aging.

## Background

Oocyte quality is an important factor in the success of animal development. Oocyte quality decreases with aging, thereby increasing errors in fertilization, chromosome segregation, and embryonic cleavage [1-3]. However, the mechanisms underlying the age-related decrease in oocyte quality remain incompletely understood.

*Caenorhabditis elegans* is a leading model for studying aging because of its short lifespan (∼2 to 3 weeks) and the conservation of longevity pathways from *C. elegans* to humans [4]. In particular, *C. elegans* has been developed as a model for studying age-related decline in fertility [5]. Mutant analyses using *C. elegans* have revealed various genes and signaling pathways that affect aging [6-8]. The molecular processes involved in the age-related regulation of oocyte quality are shared between *C. elegans* and mammals [9].

In *C. elegans* hermaphrodites, sperm are produced during the larval stage and stored in the spermatheca; oocytes are produced continually during the adult stage. Mature oocytes are transported to the spermatheca for fertilization, and the resulting embryos are pushed into the uterus and then laid through the vulva (Fig. 1a). In young animals, almost all transported oocytes are fertilized and almost all fertilized eggs are viable whereas older animals produce a significant number of unfertilized oocytes and inviable eggs, suggesting that oocyte quality declines with aging [9].

**Figure 1.**
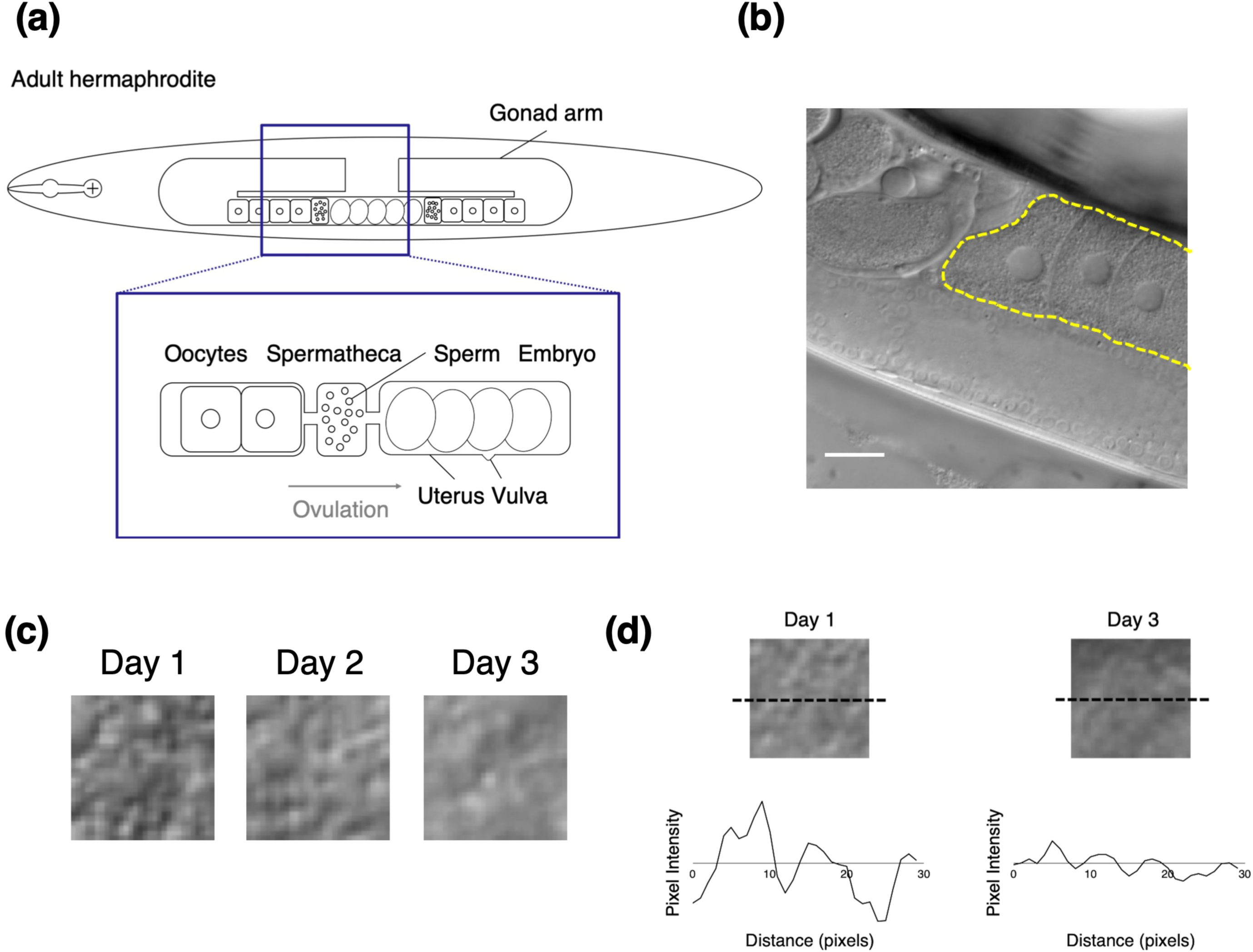
Observation of *C. elegans* oocytes by using Nomarski DIC microscopy. (a) Schematic representation of an adult hermaphroditic gonad. Oocytes mature and enter the spermatheca, where they are fertilized by the accumulated sperm. Embryos are pushed into the uterus and then laid through the vulva. (b) Representative DIC image of oocytes in gonad. The dotted yellow line surrounds the oocytes. Scale bar, 20 μm. (c) Examples of extracted images of the cytoplasmic texture in oocytes from 1-, 2-, and 3-day-old adults. (d) Cytoplasmic texture images and profile plots of Day 1 and Day 3 oocytes. The dotted black line indicates the position of the horizontal profile plot.

Age-related changes in oocytes are found not only in function but also in appearance [3]. In *C. elegans*, aged oocytes shrink, the contacts between oocytes become loose, and oocytes fuse into large clusters [5,9]. However, there is a lack of methods to objectively quantify age-related changes in oocyte appearance; such a method would provide information for clarifying the relationship between changes in the quality of oocytes and their appearance with aging.

Image processing of cell appearance has been applied to various branches of biomedical research, including the identification of malignant cells and detection of cancer [10,11], analysis of morphological changes [12], and the classification of cell populations with different functions [13,14]. Texture analysis is one method of classifying biomedical images [15]. In addition, modern machine learning methods, such as deep learning, have recently been applied to various biological applications [16]. The analysis of cell appearance by using image processing is an effective method for characterizing and classifying the status of cells.

The Gray-level Co-occurrence Matrix (GLCM) is a well-known statistical method for examining textures and is widely used to describe spatial properties [17]. The GLCM approach has been used in various biomedical applications, including cell recognition, evaluation of ultrastructural changes, and textural classification of medical images [12,18-20]. For an image with G gray levels, the GLCM is an estimate of the second-order joint probability *P*(*i, j* | *d, θ*) of two pixels with gray levels *i* and *j* (0 ≤ *i* < G, 0 ≤ *j* < G) that are *d* pixels apart from each other along direction *θ*.

To objectively describe changes in the appearance in oocytes with aging, we used Nomarski differential interference contrast (DIC) microscopy to view and characterize *C. elegans* oocytes. Nomarski DIC microscopes produce contrast by visually displaying the optical phase gradient. DIC microscopy can capture images of transparent objects without chemical staining and is widely used to observe nuclei, nucleoli, and granular structures within *C. elegans* cells [21,22]. We focused on the cytoplasmic texture because it would reflect the internal status of oocytes more directly than does morphologic appearance, such as the size or shape of oocytes. To quantify the age-associated changes in the cytoplasmic texture of *C. elegans* oocytes, we propose the image feature “Max-min Value” (Mm Value) for measuring textural roughness. We thus use Mm Value to reveal quantitative differences between young and aged oocytes. In addition, we apply methods based on GLCM, a second-order statistical method of texture analysis, to investigate differences in features between young and aged oocytes.

## Results

### Mm Value was characteristic of the age-associated change in the cytoplasmic texture of *C. elegans* oocytes

To quantify age-associated changes in oocyte appearance, we used Nomarski DIC microscopy to observe the oocytes of 1-, 2-, and 3-day-old *C. elegans* adults (hereafter called Day 1, Day 2, and Day 3 adults, respectively). In DIC microscopic images, the nucleus appears as a smooth, round region in the center of the oocyte, and the cytoplasm is rough (Fig. 1b). As previously reported [5,9], we noted various morphologic differences (accumulation of oocytes, oocyte size, cavities, and cluster formation) between Day 1 and Day 3 adults. In addition, we found that the cytoplasmic texture was rougher in appearance on Day 1 than Day 3 (Fig. 1c). Day 1 worms showed larger changes in pixel intensity than Day 3 (Fig. 1d).

To examine whether the texture changes can be characterized quantitatively, we performed a computational texture analysis of the oocyte cytoplasm. To this end, we defined an image feature, the “Max-min Value (Mm Value),” as follows. Mm Value is calculated by a moving window operation. The maximum and minimum intensities within a moving window of W×W pixels are obtained. Then the difference between maximum and minimum intensities is calculated. Mm Value is defined as the mean of the difference calculated by applying the moving window to the entire image (Fig. 2a). In a rough texture image, the Mm Value is expected to be high, whereas in a smooth texture image, the Mm Value is expected to be low.

**Figure 2.**
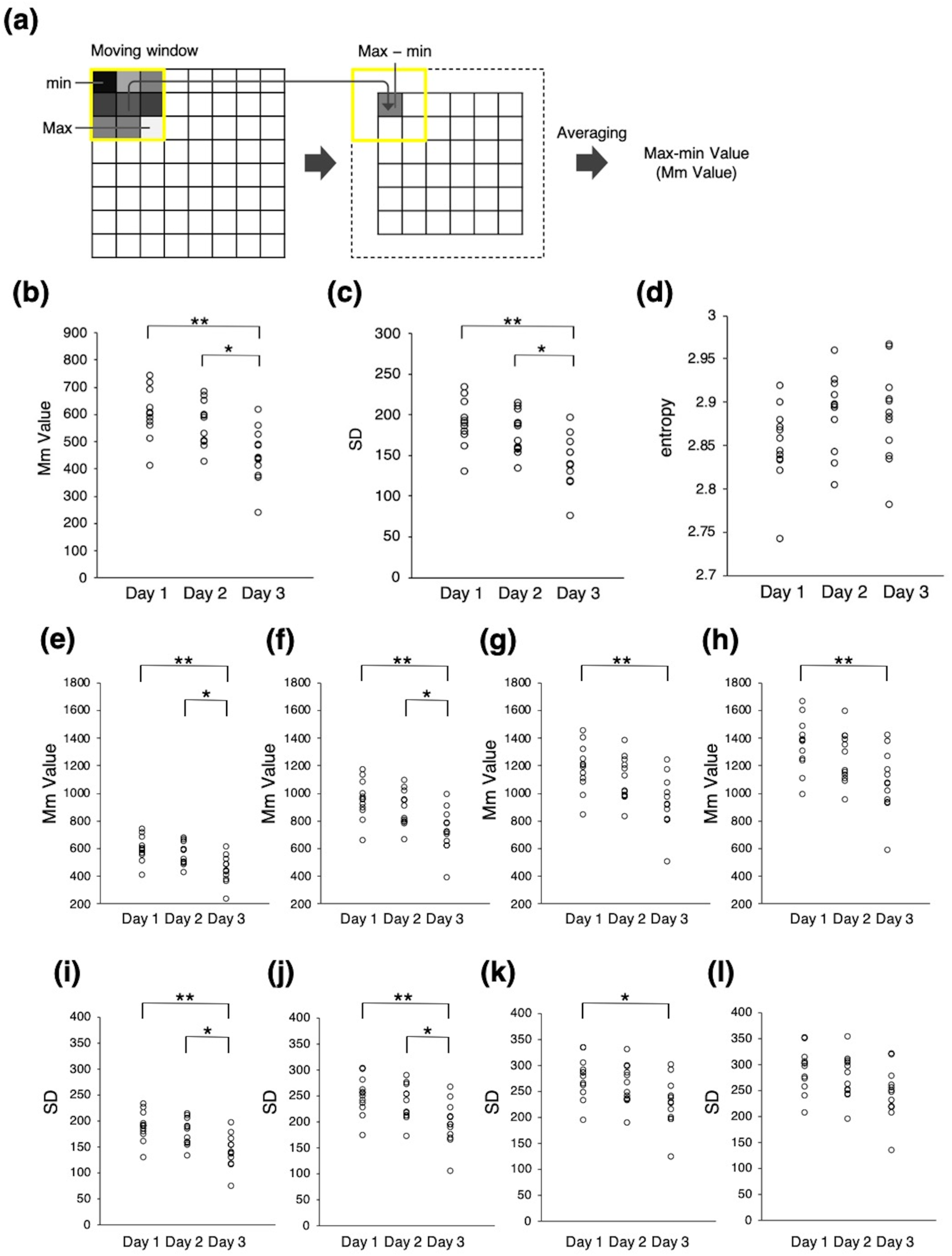
First-order statistical analysis of the age-associated change in the cytoplasmic texture of *C. elegans* oocytes. (a) Algorithm for calculating the Max-min Value (Mm Value), which is the mean of the difference between the maximum and minimum intensities within each moving window. (b–d) Comparison of the first-order statistical features of (b) Mm Value, (c) SD, and (d) entropy (3×3-pixel window) between Day 1, Day 2, and Day 3 oocytes. Circles indicate individual animals (n = 12 animals in each age group, pooled from two experiments). Asterisks indicate statistical significance (\**P* < 0.05, \*\**P* < 0.01; Tukey–Kramer test). (e–l) Comparison of (e–h) Mm Values and (i–l) SD between Day 1, Day 2, and Day 3 oocytes by using (e and i) 3×3, (f and j) 5×5, (g and k) 7×7, or (h and l) 9×9 pixels per window. Circles indicate individual animals (n = 12 animals in each age group, pooled from two experiments). Asterisks indicate statistical significance (\**P* < 0.05, \*\**P* < 0.01; Tukey– Kramer test).

When we calculated the Mm Value of the cytoplasmic texture in Day 1, Day 2, and Day 3 oocytes, we found that the Day 3 Mm Value was significantly smaller than that for Days 1 and 2 when a moving window of 3×3 pixels was used (n = 12 animals in each age group; Fig. 2b and Additional file 1: Fig. S1). Mm Values did not differ significantly between Days 1 and 2. These results suggest that the Mm Value decreases with aging and can be used to quantitatively characterize the age-associated change in the cytoplasmic texture of *C. elegans* oocytes.

To examine whether general first-order statistics quantitatively characterize the age-associated change in cytoplasmic texture, we calculated “SD” and “entropy”, which are the mean SD and entropy of local pixel intensities in a moving window. We set the size of the moving window to 3×3 pixels. Similar to the Mm Value, the SD was smaller on Day 3 than Days 1 and 2 and did not differ significantly between Days 1 and 2 (n = 12 animals in each age group; Fig. 2c). In contrast, entropy was similar among all 3 age groups (n = 12 animals in each age group; Fig. 2d). Therefore the change in cytoplasmic texture in aging *C. elegans* oocytes can be characterized quantitatively by using the Mm Value or SD but not entropy.

To compare the performance of Mm Value and SD, we set the size of the moving window to 3×3, 5×5, 7×7, and 9×9 pixels and then recalculated both parameters (Fig. 2e–l; Additional file 2: Fig. S2 shows the Mm Value at even larger window sizes [maximum, 29×29 pixels]). When a moving window of 3×3, 5×5, or 7×7 pixels was used, the Mm Value and SD of Day 1 were larger than those of Day 3. However, at a window size of 9×9 pixels, the Mm Value of Day 1 remained larger than that of Day 3, but SD did not differ between Day 1 and Day 3 (Fig. 2h and l). In addition, the Mm Values were statistically different between Day 1 and Day 3 for window sizes of 3×3 to 17×17 but not for larger windows (Additional file 2: Fig. S2). These results suggest that, compared with SD, Mm Value more robustly characterized the age-associated change in cytoplasmic texture.

### The second-order statistic GLCM varied with the age-associated change in cytoplasmic texture in *C. elegans* oocytes

GLCM is an estimate of the second-order joint probability *P*(*i, j* | *d, θ*) that two pixels with gray levels *i* and *j* are *d* pixels apart from each other in the direction *θ* (Fig. 3a). To examine whether a second-order statistic more significantly characterizes age-associated texture changes than Mm Value, we used the GLCM-based texture feature *Correlation* (COR), which is a measurement of the gray level linear dependencies of pixels at specified positions relative to each other. We calculated COR of the cytoplasmic texture of Day 1, Day 2, and Day 3 oocytes.

**Figure 3.**
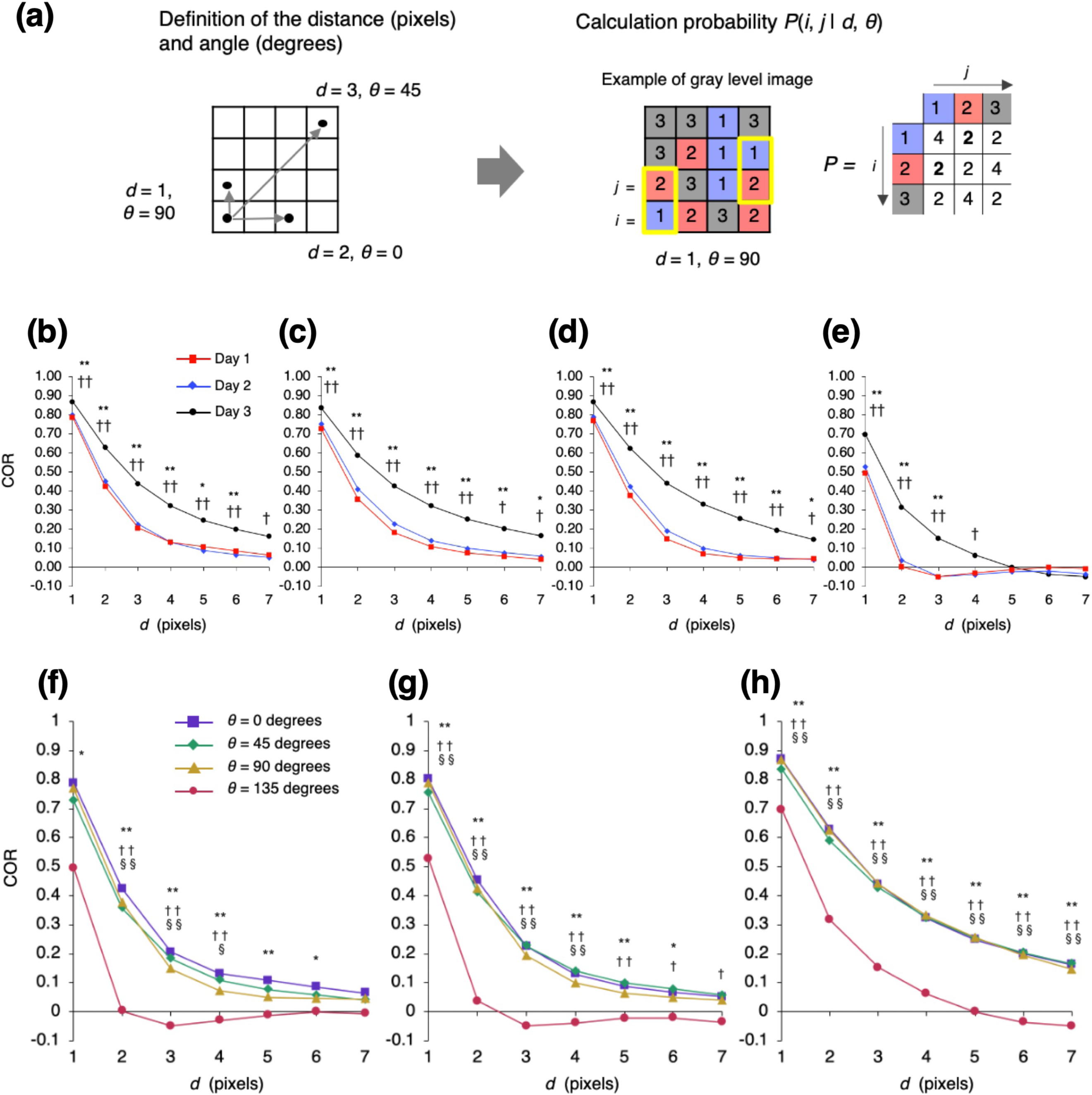
Second-order statistical analysis of the age-associated change in the cytoplasmic texture of *C. elegans* oocytes. (a) Algorithm for calculating the Gray-Level Co-Occurrence Matrix (GLCM). First, we define a spatial relationship by using the parameters distance *d* and angle *θ*. We then calculate the second-order joint probability *P*(*i, j* | *d, θ*) of two pixels with gray levels *i* and *j* (0 ≤ *i* < G, 0 ≤ *j* < G). To calculate *P*(*i, j* | *d, θ*), we sum the number of pixels with paired intensities (*i* and *j*) in the defined spatial relationship. For example, when the distance is 1 pixel and the direction is 90 degrees, the calculated number of pixels with *i* = 1 and *j* = 2 or *i* = 2 and *j* = 1 is 2. The co-occurrence matrix defined is symmetric. (b–e) The *Correlation* (COR) curve as a function of distance *d*. Here, *θ* is set to (b) 0, (c) 45, (d) 90, and (e) 135 degrees. Comparison of the mean COR of cytoplasmic texture between Day 1, Day 2, and Day 3 oocytes (n = 12 animals in each age group, pooled from two experiments). Symbols indicate statistical significance (Tukey–Kramer test) between Day 1 and Day 3 oocytes (\**P* < 0.05, \*\**P* < 0.01) or Day 2 and Day 3 oocytes (†*P* < 0.05, ††*P* < 0.01). (f–h) COR as a function of distance *d*. COR of cytoplasmic texture in (f) Day 1, (g) Day 2, and (h) Day 3 oocytes. Comparison of the mean COR between *θ* of 0, 45, 90, and 135 degrees (n = 12 animals in each angle group, pooled from two experiments). Symbols indicate statistical significance (Tukey–Kramer test) between 0 and 135 degrees (\**P* < 0.05, \*\**P* < 0.01), 45 and 135 degrees (†*P* < 0.05, ††*P* < 0.01), or 90 and 135 degrees (§*P* < 0.05, §§*P* < 0.01).

COR decreased as *d* got larger and converged to around zero (Fig. 3b–e). For various *θ* values, the *d* value at which COR converged was larger in Day 3 oocytes than in Day 1 or 2 oocytes; for example, in the case where *θ* = 135, COR in Day 3 oocytes converged to zero when *d* = 5, whereas COR in Day 1 or Day 2 oocytes converged to zero when *d* = 2 (Fig. 3e). We found that, at several levels of parameters *d* and *θ*, COR was significantly larger in Day 3 oocytes than in Day 1 and Day 2 oocytes. In particular, COR in Day 3 oocytes was most significantly larger than that in Day 1 when *d* = 1 and *θ* = 135 (*P* = 1.0×10^−8^; n = 12 animals in each age group; Tukey–Kramer test on COR). In particular, the *P* value for this parameter set was four orders of magnitude lower than the lowest *P* value obtained by using Mm Value (window size, 3×3 pixels; *P* = 6.0×10^−4^; n = 12 animals in each age group; Tukey–Kramer test on Mm Value). These results suggest that COR effectively characterized the age-associated changes in cytoplasmic texture. COR was able to more significantly characterize the differences between Day 1 and Day 3 than the Mm Value when using an appropriate parameter set. In addition to COR, we tested several texture-associated features based on GLCM, including *Angular Second Moment* (ASM), *Contrast* (CON), *Inverse Difference Moment* (IDM), and *Entropy* (ENT), but COR continued to yield the best characterization (Additional file 3: Fig. S3).

In general, for all oocytes regardless of age, COR at *θ* = 135 degrees was the smallest among all angles tested (Fig. 3f– h).

### Sample orientation does not influence the angle dependency of cytoplasmic texture

Images from DIC microscopy have a shadow-cast appearance oriented in the shear direction of the prism^23^. The texture characterized by using COR was angle-dependent, i.e., COR at *θ* = 135 was smaller than at any other angle (Fig. 3f–h). To examine whether this angle dependency is due to either the orientation of the worms or the imaging system itself, we calculated the COR after rotating worm samples to 0, 45, 90, or 135 degrees relative to horizontal (i.e., 0 degrees) (Fig. 4a–d). If sample orientation causes the angle dependency, then the angle-dependent property of the COR should change depending on sample orientation. Conversely, if the angle dependency is due to the imaging system itself, then orientation should have no effect on this property. We found that—regardless of sample orientation—COR at *θ* = 135 was smaller than at other angles (n = 12 animals in each angle group; Fig. 4e–h). This result indicates that the angle-dependent property of oocyte cytoplasmic texture is due to the imaging system itself, rather than to the orientation of the worm imaged.

**Figure 4.**
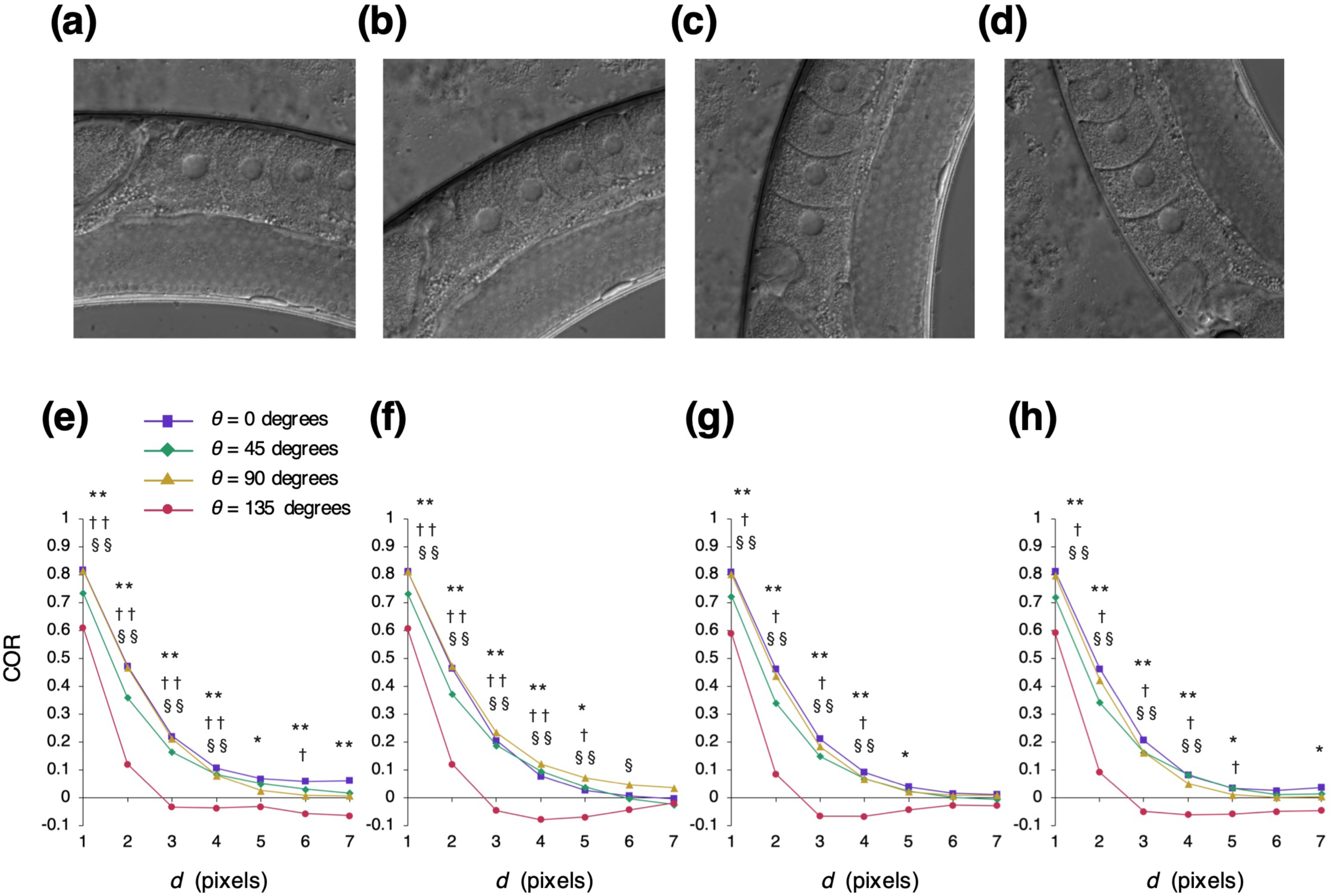
Comparison of COR between angle parameters by using rotated worms. (a–d) Representative images of worms rotated (a) 0, (b) 45, (c) 90, or (d) 135 degrees from horizontal (i.e., 0 degrees). (e–h) The COR curve as a function of distance *d*. The COR of worms rotated (e) 0, (f) 45, (g) 90, and (h) 135 degrees from horizontal. Comparison of the mean of COR between *θ* of 0, 45, 90, and 135 degrees (n = 10 animals in each angle group). Symbols indicate statistical significance (Tukey–Kramer test) between 0 and 135 degrees (\**P* < 0.05, \*\**P* < 0.01), 45 and 135 degrees (†*P* < 0.05, ††*P* < 0.01), or 90 and 135 degrees (§*P* < 0.05, §§*P* < 0.01).

### Changing smoothness or simulating large structure cannot reproduce the age-associated changes in cytoplasmic texture

To elucidate the factor that causes the age-associated texture change that is characterized by Mm Value or COR, we visually compared Day 1 and Day 3 oocytes. We considered two factors that might underlie difference in texture with aging: cytoplasmic smoothness (that is, Day 3 cytoplasm appeared smoother than Day 1) and the distribution of large structures (that is, large structures were distributed irregularly in the cytoplasm of Day 3 oocytes but were homogenous throughout Day 1 cytoplasm) (Fig. 5a).

**Figure 5.**
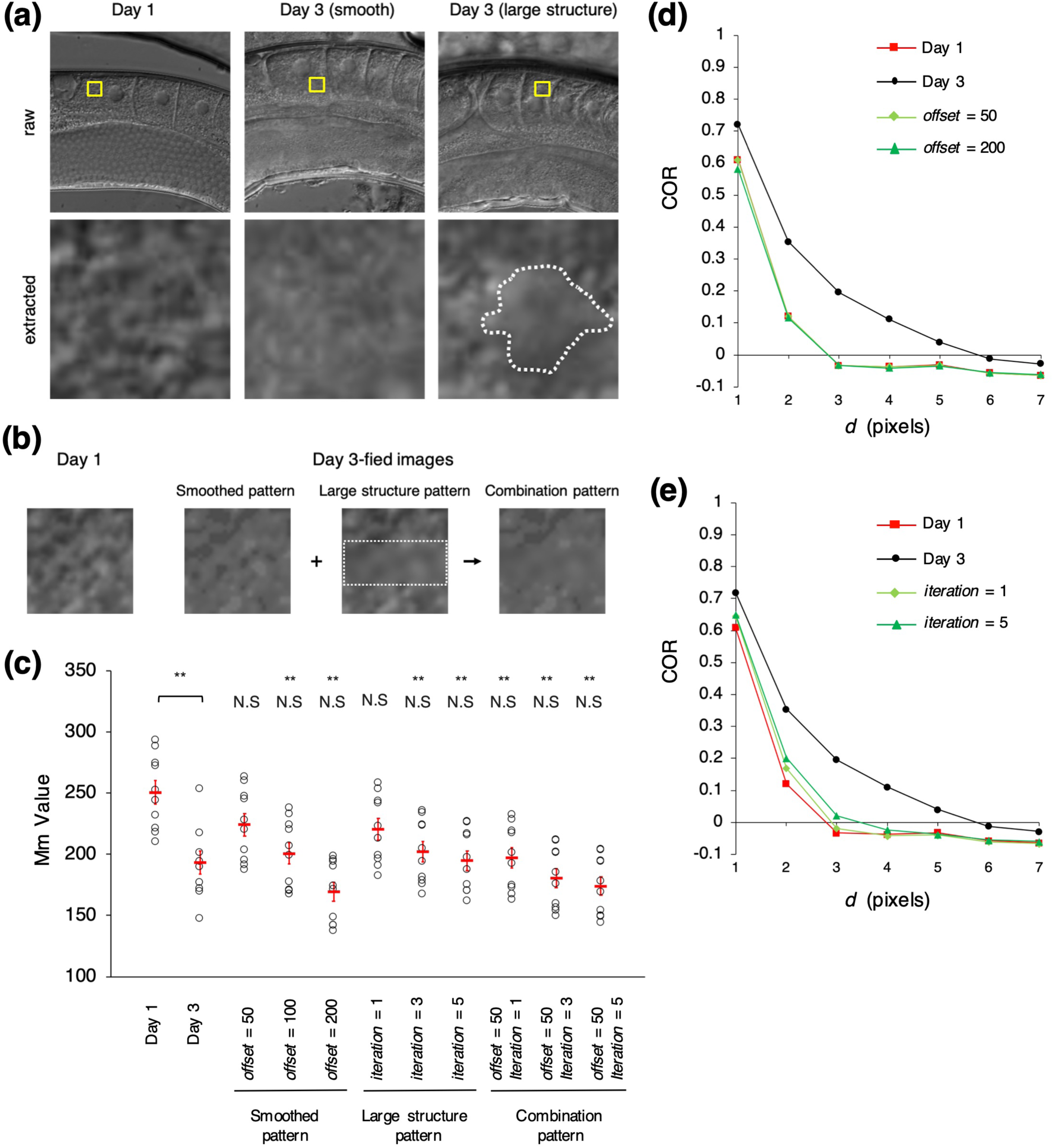
Day 3-fied images created by using Day 1 images. (a) Comparison of cytoplasmic texture between Day 1 and Day 3 oocytes. Top, oocyte images; the yellow area indicates the extracted area. Bottom, extracted images. The Day 3 sample (middle) appears to have a smoother texture than the Day 1 oocyte (left). The Day 3 oocyte (right) contains large structures (dotted white line), whereas the Day1 sample is homogeneous. (b) Day 1 image and three Day 3-fied images (left, Smoothed pattern; middle, Large Structure pattern; right, Combination pattern) based on the Day 1 image. The dotted white line indicates the filtered area. (c) Mm Value of actual Day1, actual Day 3, and three Day 3-fied images. Comparison of Mm Value between Day 1 and Day 3-fied images or actual Day 3 and Day 3-fied images. The Smoothed pattern parameter *offset* was set to 50, 100, and 200. The Large Structure pattern parameter *iteration* was set to 1, 3, and 5. The Combination pattern parameters (*offset, iteration*) were set to (50, 1), (50, 3), and (50, 5). Circles indicate the Mm Values of individual animals (n = 10 animals in each age group; orientation of the worms is 0 degrees); red bars indicate the mean values. Error bars indicate SEM. Asterisks indicate statistical significance (Tukey–Kramer test) between actual Day1 and actual Day 3 or actual Day 1 and Day 3-fied images (\*\**P* < 0.01); N.S., no significant difference between actual Day 3 and Day 3-fied images. (d and e) COR as a function of distance *d* at *θ* = 135 degrees. The mean COR of the cytoplasmic texture in actual Day 1, actual Day 3, and Day 3-fied oocyte images (n = 10 animals in each age group; orientation of the worms is 0 degrees) created due to (d) Smoothed pattern (*offset* = 50 or 200) or (e) Large Structure pattern (*iteration* = 1 or 5).

To examine whether these factors caused the texture changes, we created ‘Day 3-fied images’ by applying image processing to Day 1 images. We created three patterns of Day 3-fied images by manipulating the factors. The first pattern, ‘Smoothed pattern’, was created by smoothing the images of Day 1. That is, the maximum and minimum intensities of Smoothed pattern images were normalized to minimum + *offset* and maximum − *offset* by using the minimum and maximum intensities from Day 1 oocytes. The second pattern, ‘Large Structure Pattern’, was generated by applying a Gaussian filter at *iteration* times to the center part of the Day 1 images. We created the third pattern, ‘Combination Pattern’, by first smoothing the Day 1 images and then generating large structures on them (Fig. 5b).

We then compared Mm Value and COR between Day 3-fied, actual Day 1, and actual Day 3 oocyte images (n = 10 animals in each age group; Fig. 5c). We set the parameter *offset* in the Smoothed pattern to 50, 100, or 200 and the parameter *iteration* in the Large Structure pattern to 1, 3, or 5. If the properties of the Day 3-fied images are similar to the actual images, the features of the Day 3-fied images will be similar to those of actual Day 3 images and differ from those of actual Day 1 images. For Smoothed pattern images, Mm Value for the Day 3-fied image did not differ significantly from Day 3 data but was significantly smaller than that of Day 1 (Fig. 5c; *offset* = 100, 200). The Large Structure and Combination patterns yielded similar findings as for the Smoothed pattern (Fig. 5c). In other words, all patterns of the Day 3-fied images reproduced the difference in Mm Value between Day 1 and Day 3 oocytes and were similar to actual Day 3 oocytes.

Next, we calculated COR for Day 3-fied, actual Day 1, and actual Day 3 images (n = 10 animals in each age group). For the actual images, COR converged to zero at larger *d* values for Day 3 than Day 1 (Fig. 3b–e). However, COR of Day 3-fied images converged to zero at the almost same *d* as Day 1 for both the Smoothed and Large Structure patterns (Fig. 5d and e) as well as the Combination Pattern (Additional file 4: Fig. S4). None of the three patterns of Day 3-fied images accurately represented the difference in COR between actual Day 1 and Day 3 images. This finding suggests that the age-associated change in cytoplasmic texture cannot be artificially reproduced by manipulating the smoothness of the image or adding irregular large structures.

### Synthetic images with different sizes of granules recapitulated the difference in COR between Day 1 and Day 3 oocytes

The Day 3-fied images created by using the Day 1 images failed to reproduce the difference in COR between Day 1 and Day 3 images, i.e., COR converged to zero at larger *d* in Day 3 than Day 1 images (Fig. 5d and e). To investigate what factor causes the difference in the convergence of COR between Day 1 and Day 3 oocytes, we created simple synthetic images that were based on two hypotheses: that the (1) number or (2) size of granules in the cytoplasm changes with aging. We therefore evaluated the convergence of COR in the synthetic images by varying granule number (*N*) or size (*R* pixels) (Fig. 6a).

**Figure 6.**
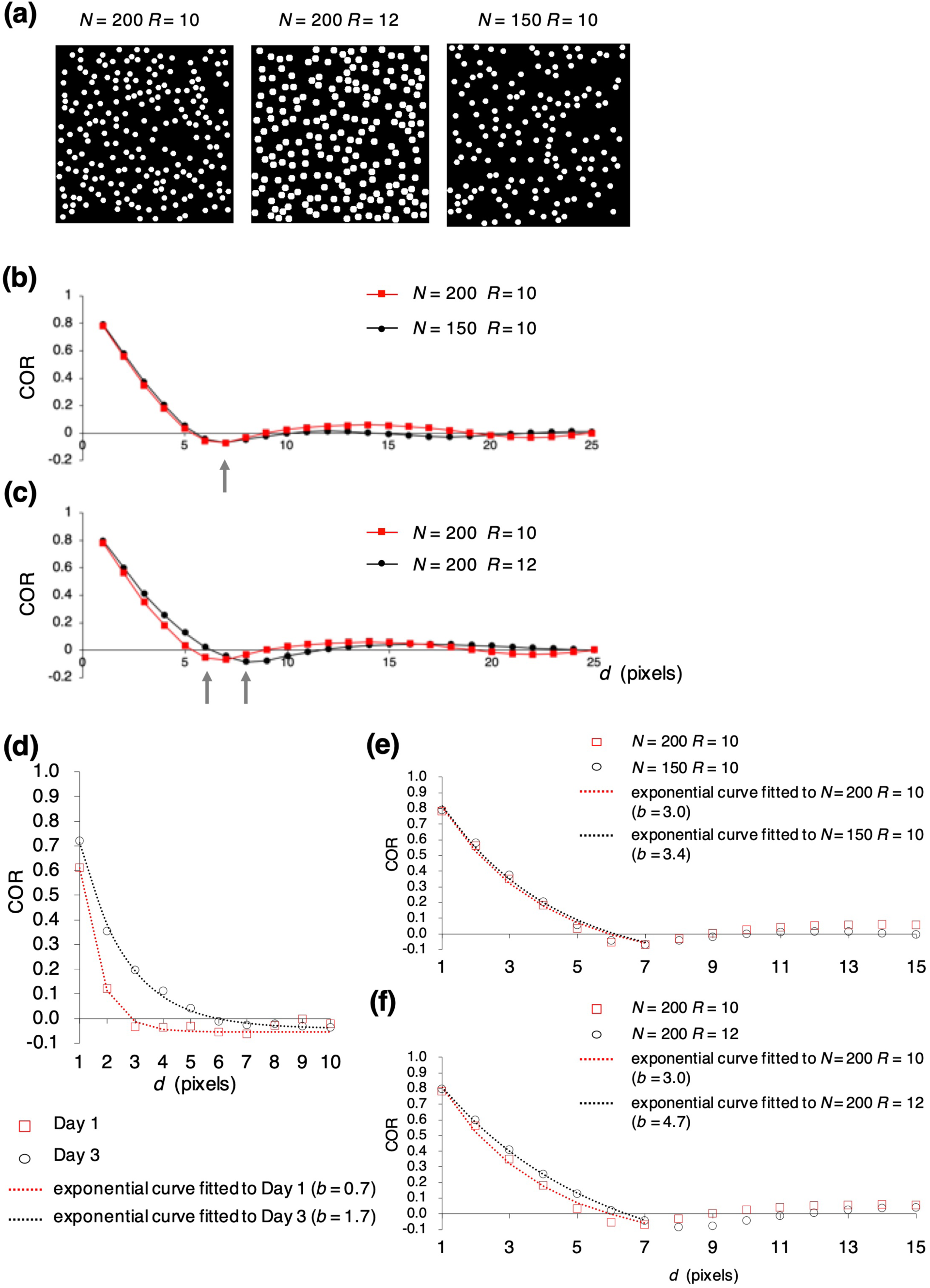
Synthetic images created by using different numbers and sizes of particles. (a) Synthetic images created by setting the number (*N*) and size (*R* pixels) of particles to (left) 200 and 10, (middle) 200 and 12, or (right) 150 and 10, respectively. (b and c) COR as a function of distance *d* at *θ* = 135 degrees. Comparison of COR properties after changing the (b) number (*N*) or (c) size (*R* pixels) of particles. Gray arrows indicate *d* converging to zero. (d) The mean COR at *θ* = 135 degrees in Day 1 and Day 3 oocytes (n = 10 animals in each age group; orientation of the worms is 0 degrees) and the curves fitted to each set of COR data. (e and f) The COR of the synthetic images and the curves fitted to each set of COR data. Comparison of the properties of the fitted curves due to changing particle (e) number (*N*) or (f) size (*R* pixels).

When we varied granule number but kept the granule size constant at *R* = 10, COR in the synthetic images converged at approximately the same *d* regardless of whether 200 or 150 granules were present (Fig. 6b). In contrast, when we varied the granule size but kept the granule number constant at *N* = 200, COR converged at a larger *d* when the granule size was 12 pixels compared with 10 pixels (Fig. 6c). Therefore, merely altering the number of the granules did not recapitulate the difference in the convergence of COR between Day 1 and Day 3 oocytes. These results suggest that changing the size of cytoplasmic granules can reproduce the difference in the convergence of COR between actual Day 1 and Day 3 oocytes.

To objectively assess whether changing granule size in synthetic images yields the anticipated difference in COR, we exponentially approximated COR curves by using Equation (1), where *a* is the amplitude, *b* is the distance until convergence to zero, and *c* is the offset.

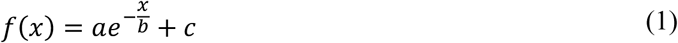

In the approximation function, as *b* increases, the *d* value at which *f*(*x*) converged to zero increases. The difference in the *d* at which COR converges to zero between Day 1 and Day 3 should reflect the difference in *b* values. When we compared the *b* in the function approximating COR between Day 1 and Day 3, the *b* of Day 3 (*b* = 1.7) was larger than that of Day 1 (*b* = 0.7; Fig. 6d). Next, we approximated COR of the synthetic images. The difference in *b* was greater when we manipulated granule size *R* (Fig. 6f; [*N, R*] = [200, 10], *b* = 3.0; [*N, R*] = [200, 12], *b* = 4.7) than when we varied granule number (*N*) (Fig. 6e; [*N, R*] = [200, 10], *b* = 3.0; [*N, R*] = [150, 10], *b* = 3.4). The *b* was larger for the larger granule size than the smaller granule size. The results objectively suggest that the difference of the COR property between Day 1 and Day 3 is able to be represented by changing the size of granules.

### Granules in *C. elegans* oocytes are larger on Day 3 than Day 1

In the synthetic images, changing the granule size reproduced the difference in COR between Day 1 and Day 3 oocytes, thus suggesting that cytoplasmic granules in *C. elegans* oocytes might change in size with aging. To examine whether granules in DIC images demonstrated age-associated size variation, we manually measured granules and compared their size on Days 1 and 3 (n = 8 animals in each age group Fig. 7a). Granules were significantly larger in Day 3 oocytes than Day 1 oocytes (Fig. 7b).

**Figure 7.**
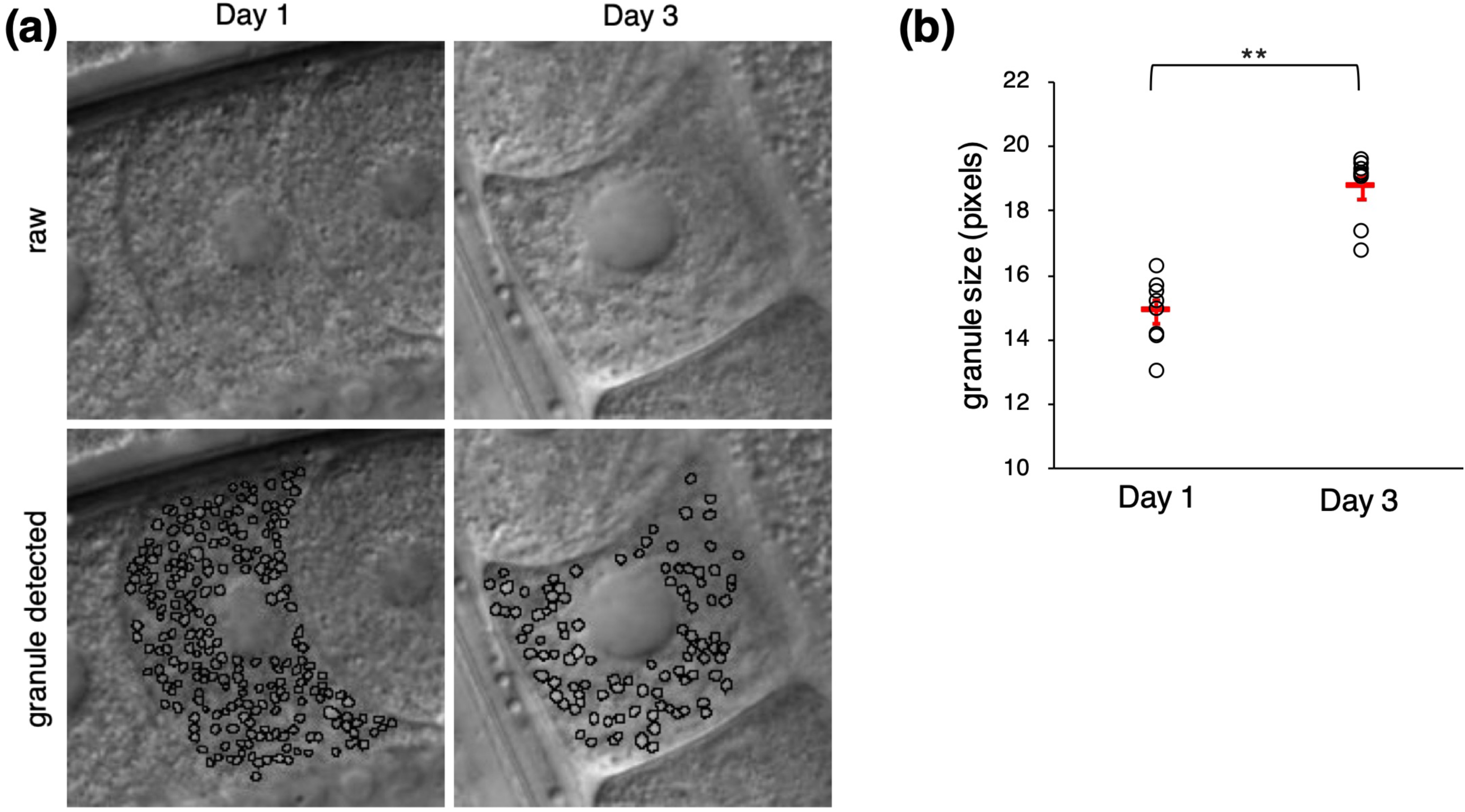
Measurement of the size of cytoplasmic granules in DIC images. (a) Representative DIC images of oocytes. Left, Day 1 oocyte. Right, Day 3 oocyte. Top, Raw images. Bottom, Manual measuring of granule areas on raw images. Many of the elongated and clustered granules in Day 3 oocytes were not measured. (b) Comparison of the size of granules manually measured in Day 1 and Day 3 oocytes. Circles indicate individual animals (n = 8 animals in each age group); red bars indicate the mean value. Error bars indicate SEM. Asterisks indicate statistical significance (\*\**P* < 0.01; Welch’s two-tailed *t* test).

## Discussion

In this study, we found that the texture of the cytoplasm in *C. elegans* oocytes varies with their age. This change was characterized quantitatively through the DIC image features of Mm Value and COR, which is based on the second-order statistic GLCM. In addition, at appropriate parameter sets, COR characterized this age-associated feature more significantly than Mm Value. Furthermore, analysis of synthetic images and measurement of the size of cytoplasmic granules suggested that the cytoplasmic granules in *C. elegans* oocytes become larger with aging.

Mm Value is a contrast feature that is calculated as the mean of the difference between the maximum and minimum intensities within successive moving windows. The statistical significance of the difference in Mm Values between Day 1 and Day 3 oocytes decreased as the window size increased (Additional file 2: Fig. S2). If the texture contrast is uniform within an oocyte, calculating Mm Value by using a window size that is smaller than the granule size enables the Mm Value to fluctuate depending on the size or density of granules. However, this variation cannot occur when the window size used for determining Mm Value might contain multiple granules, such as 13×13, 15×15 or 17×17. In this case, the Mm Value for Day 1 oocytes remained significantly larger than for Day 3 (Additional file 2: Fig. S2). Therefore, texture contrast might decrease with aging. However, the utility of Mm Value for characterizing the age-associated change in cytoplasmic texture disappeared when window size was set to 19×19 pixels or larger. When determined by using windows sufficiently large to contain multiple granules, the significance of differences in Mm Values characterizing age-associated differences in texture decreased as window size increased. These findings may indicate that texture contrast changes in a spatially inhomogeneous manner.

Regardless of window size, the Mm Values of the Day 3-fied images based on the Smoothed pattern were significantly smaller than those of Day 1 (Additional file 5: Fig. S5). In addition, the Mm Values of the Day 3 images did not differ from those of Day 1 images when the window was larger than 23×23. Furthermore, contrast in the Smoothed pattern image was decreased due to normalization to the overall texture throughout the image. Therefore, the results suggest that texture contrast does not change homogeneously from Day 1 to Day 3. The inhomogeneous changes of the texture contrast may reflect the relationship between the Mm Values of Day 1 and Day 3. Given that changes in the optical phase gradient can alter contrast in DIC images, the age-associated decrease of contrast might reflect a change in granule content due to chemical modification or a difference in content quantity. Similar to the Mm Values of Day 3 images, the Mm Values of Day 3-fied images that were based on the Large Structure and Combination patterns provided fewer significant difference from Day 1 images as window size increased. (Additional file 5: Fig. S5). The Day 3-fied images based on the Large Structure and Combination patterns were created by changing the contrast of Day 1 spatially inhomogeneously.

COR significantly characterized age-associated variations when using smaller *d* but not when using larger *d* (Fig. 3b-e). That COR was more effective at characterizing age-associated differences at small *d* values suggests that some small textural feature changes with aging. Therefore, COR likely significantly characterized age-associated changes in texture by recognizing small structures in the cytoplasm.

The convergence of COR in the Day 3-fied images differed markedly from those of the actual Day 3 (Fig. 5d and e). The synthetic images revealed the effect of granule size—rather than number—on the difference in COR between Day 1 and Day 3 (Fig. 6b and c). These results suggest that the size of granules is the major factor affecting the convergence of COR. In addition, we found that cytoplasmic granules in the Day 3 oocytes tended to be larger than those in the Day 1 oocytes (Fig. 7b), suggesting that cytoplasmic granules in *C. elegans* oocytes become larger with age.

At the optimal parameter setting (*d* = 1, *θ* = 135) and a window of 3×3 pixels, COR characterized cytoplasmic texture more significantly than the Mm Value. However, the COR-based characterization is not always more significant than that of the Mm Value. The effectiveness of COR depends on the angle *θ*. Using the images of the rotated worms suggested that the age-associated differences in texture include an angle-dependent property that is intrinsic to the DIC imaging system. At appropriate parameters, COR characterized the changes in texture more significantly than Mm Value, but Mm Value was more informative at a broader range of parameters and was not particularly influenced by the angle parameter.

Reported age-associated changes in the morphologic appearance of *C. elegans* oocytes include oocyte shrinkage, loosened contacts, and aggregation into large clusters [5,9]. In addition, we have revealed age-associated changes in the cytoplasmic texture of *C. elegans* oocytes. This change could be quantitatively characterized through several image features. Cytoplasmic granules would reflect the internal status of oocytes more directly than does the external morphologic appearance. Quantitative analysis of cytoplasmic texture and measurement of granules suggest that the cytoplasmic granules in *C. elegans* oocytes become larger with aging. Various image features can be used as objective methods to quantify age-associated differences in oocyte appearance, which may reflect oocyte fertility. To use the texture of oocytes to classify them according to their age, image features should differ markedly relative to oocyte age. Although various image features characterized age-associated changes in cytoplasmic texture, their distribution overlapped between age groups, perhaps reflecting insufficiently sensitive methods. Recognizing and classifying textural changes accurately may require multi-dimensional analysis by using additional image features or the application of machine learning methods, such as deep learning. However, age-associated changes in cytoplasmic texture can be subtle: Mm Value and COR differed significantly between Day 1 and Day 3 oocytes but not between Day 1 and Day 2 oocytes.

Some of cytoplasmic granules in our DIC texture images may be yolk granules [24,25]. Yolk is a lipoprotein composed of lipids and lipid-binding proteins, called vitellogenins [26]. In *C. elegans*, vitellogenins are synthesized in the intestine and transported into maturing oocytes through endocytosis [27,28]. Yolk provides essential nutrients to the eggs to support embryonic development [28]. During reproductive senescence, the intestine continues to produce and secrete large amounts of yolk protein. In adult *C. elegans* hermaphrodites, yolk accumulates toward the end of the self-fertile reproductive period [29,30]. Provisioning of vitellogenin to embryos increases with maternal age [31] and, given that vitellogenins transport lipids into embryos [26,31], might increase the lipid content in embryos and oocytes. Together with our data, the cytoplasmic granules in aged adults might enlarge due to an increase in vitellogenin content or in vitellogenin-transported embryonic lipid content.

What is the relationship between decreased fertility and yolk accumulation with aging? Yolk accumulation may contribute to the decrease in fertility. Yolk amount appears to affect the lifespan of *C. elegans*, in that high levels of yolk appear to be detrimental and decrease lifespan [32]. In contrast, knockdown of vitellogenin expression extends lifespan [33,34]. Increased amounts of yolk might accelerate the aging of oocytes or animals, resulting in decreased fertility. Alternatively, decreased oocyte fertility might contribute to yolk accumulation. Moreover, we cannot exclude the possibility that yolk accumulation does not affect fertility directly. Further experiments are needed to clarify the relationship between yolk levels and fertility in this and other species.

## Conclusions

In this study, we found that the Mm Value and COR objectively quantify age-related changes in *C. elegans* oocyte in Nomarski DIC microscopy images. We are planning to apply these image features to publicly available Nomarski DIC microscopy images of *C. elegans* oocytes and embryos in public image databases such as Phenobank [35] and WDDD [36]. Such applications may provide clues to the molecular mechanism of oocyte aging.

## Methods

### *C. elegans* Strains and Growth Conditions

Nematodes (Bristol N2 strain) were grown under standard conditions [37]. The L4 larval stage was considered as Day 0; worms were defined as Day 1 adults at 18–24 h after L4; as Day 2 adults at 43–46 h, and as Day 3 adults at 67–70 h.

### Imaging

Worms were immobilized in polystyrene nanoparticle suspension [38] supplemented with 5-hydroxytryptamine (5-HT) [39] on agarose pads. The anterior gonad was observed by Nomarski DIC microscopy using a Leica HCX PL APO 63×/1.20 W CORR objective and an iXonX3 electron-multiplying CCD camera controlled with live cell imaging software (Andor iQ). Digital images of 512×512 pixels were converted to 14-bit TIFF format (0.25 μm per pixel). The DIC microscopy was adjusted to focus on the nucleus of the most proximal oocytes. At least 3 oocytes were imaged per animal. Images of nematodes at four angles (0, 45, 90, and 135 degrees) were obtained by rotating the Day 1 samples; 0 degrees was defined as horizontal orientation.

### Calculation of image features

To calculate image features, random regions of 30×30 pixels were extracted from the cytoplasm, without including nuclei or cell boundaries. One region was extracted from each oocyte, and the three most proximal oocytes to the spermatheca in the anterior gonad were used for each animal. Image features of individual animals were defined as the mean of those in the extracted three regions.

The first-order statistical features Mm Value, SD, and entropy were calculated by moving the local window except for the border. SD was defined as the mean of the standard deviations of the pixel intensities in the moving window. Entropy was defined as the mean of entropies (calculated according to the following the equation) in the moving window:

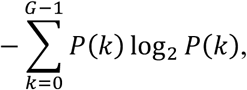

where *G* is the number of gray levels, and *P*(*k*) is the probability of occurrence of gray level *k* in the moving window.

Second-order statistical features based on GLCM—*Correlation* (COR), *Angular Second Moment* (ASM), *Contrast* (CON), *Inverse Difference Moment* (IDM), and *Entropy* (ENT)—were calculated by using the following equation and the co-occurrence matrix *P*(*i, j* | *d, θ*):

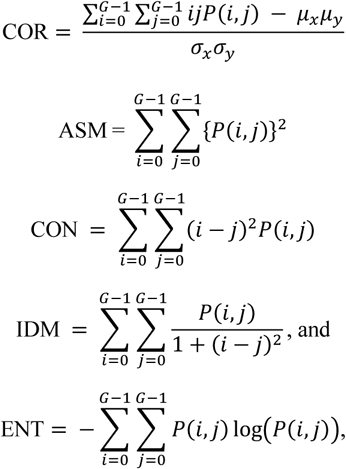

where *μ*_*x*_, *μ*_*y*_, *σ*_*x*_, and *σ*_*y*_ are the means and standard deviations in the *x* and *y* direction given by

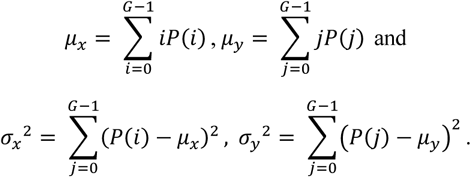

*P*(*i, j* | *d, θ*) was defined symmetric (see Fig. 3a).

### Creation of the synthetic images

The synthetic images were created by randomly locating *N* white granules with diameter *R* pixels on a black background image of 300×300 pixels. Granules did not to overlap or protrude from the border of the image.

### Fitting to the COR curves

The COR was fitted to the following exponential function through nonlinear regression analysis:

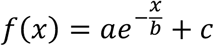

When fitting the COR of actual Day 1 and Day 3 images or synthetic images, we used the COR for *d* of 1 to 7 pixels, to approximate convergence of the properties to zero. We used data of worms oriented to 0 degrees (n = 10 animals).

### Quantification of the granule size

For each worm, we manually measured the cytoplasmic granules in the second oocyte in the spermatheca (the next most proximal oocyte). Granules were determined as non-overlapping circular or nearly circular regions where signal intensity exceeded the background intensity. Many of the elongated and clustered granules in Day 3 oocytes were not counted.

## Additional files

**Additional file 1: Figure S1.**
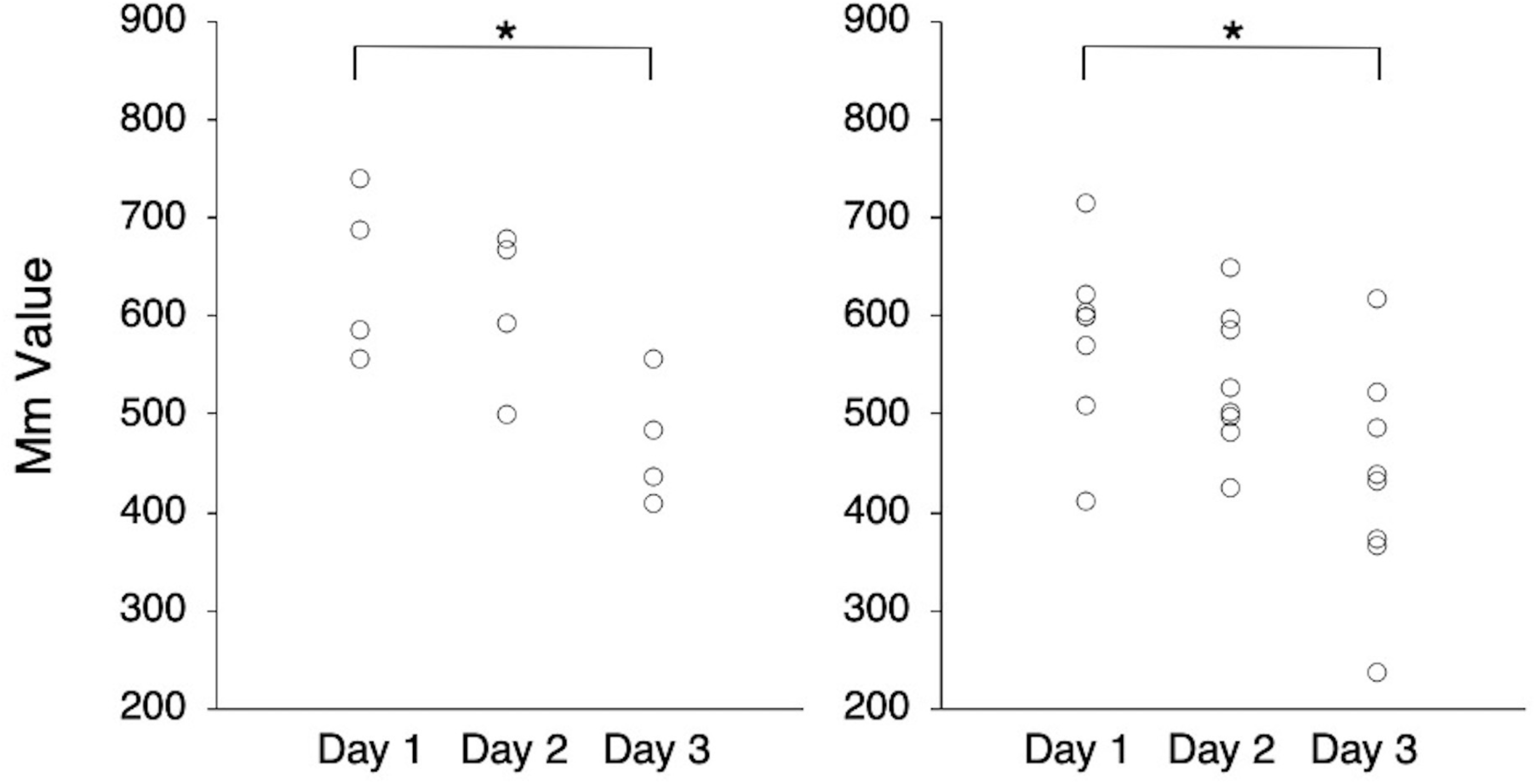
Comparison of the Max-min Value (Mm Value) in each of two experiments. Comparison of the Mm Value calculated by using a 3×3-pixel window between Day 1, Day 2, and Day 3 oocytes. Circles indicate individual animals (n = 4 or 8 animals each age group). Asterisks indicate statistical significance (\**P* < 0.05; Tukey–Kramer test).

**Additional file 2: Figure S2.**
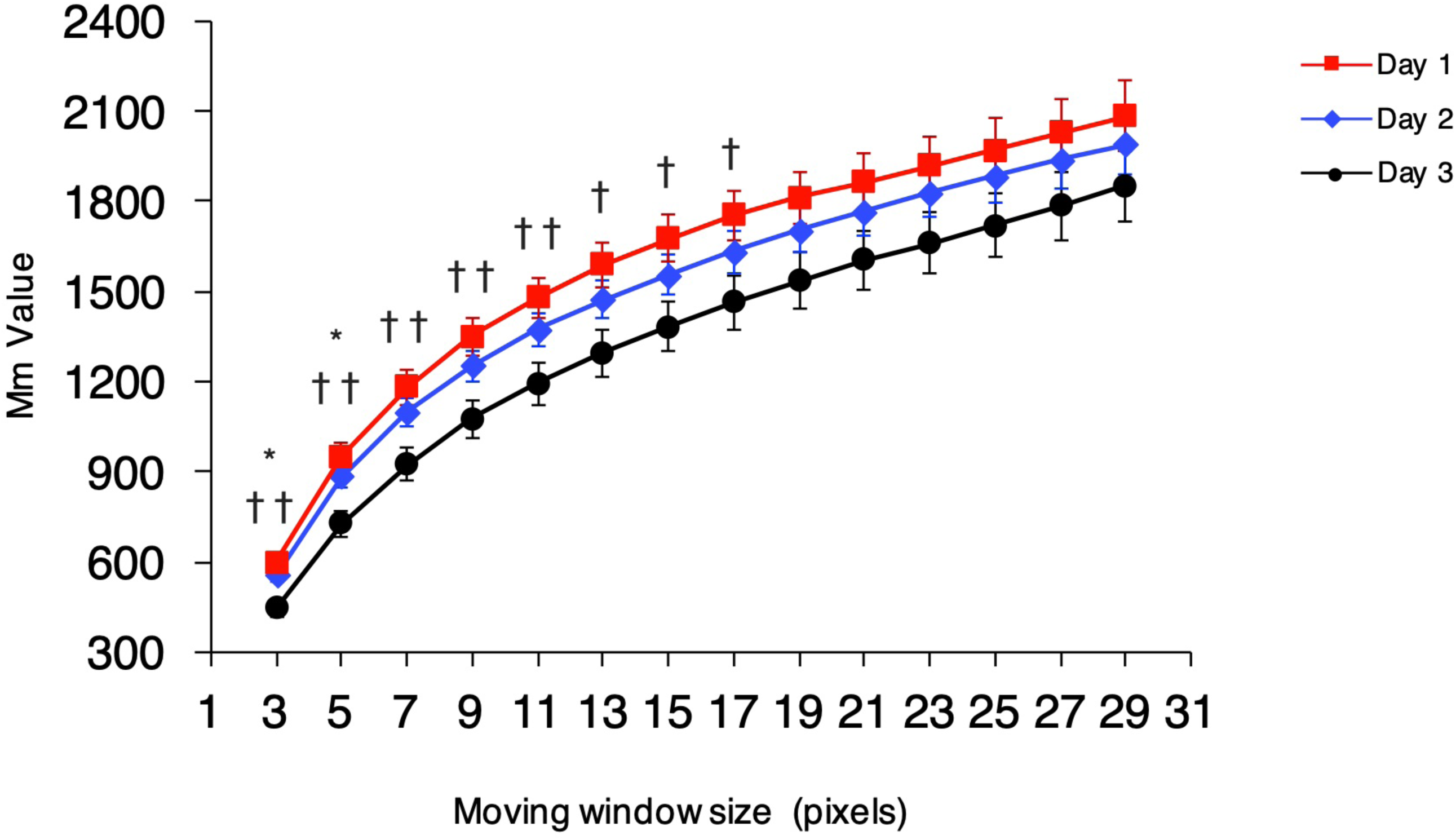
Comparison of Mm Value between oocytes from 1-, 2-, and 3-day-old adults (Day 1, Day 2, and Day 3, respectively) after changing the window size. Comparison of Mm Value between Day 1, Day 2, and Day 3 oocytes (n = 12 animals each age group, pooled from two experiments) after changing the window size from 3×3 to 29×29. Error bars indicate SEM. Symbols indicate statistical significance (Tukey–Kramer test) between Day 1 and Day 3 (\**P* < 0.05, \*\**P* < 0.01) or Day 2 and Day 3 (†*P* < 0.05).

**Additional file 3: Figure S3.**
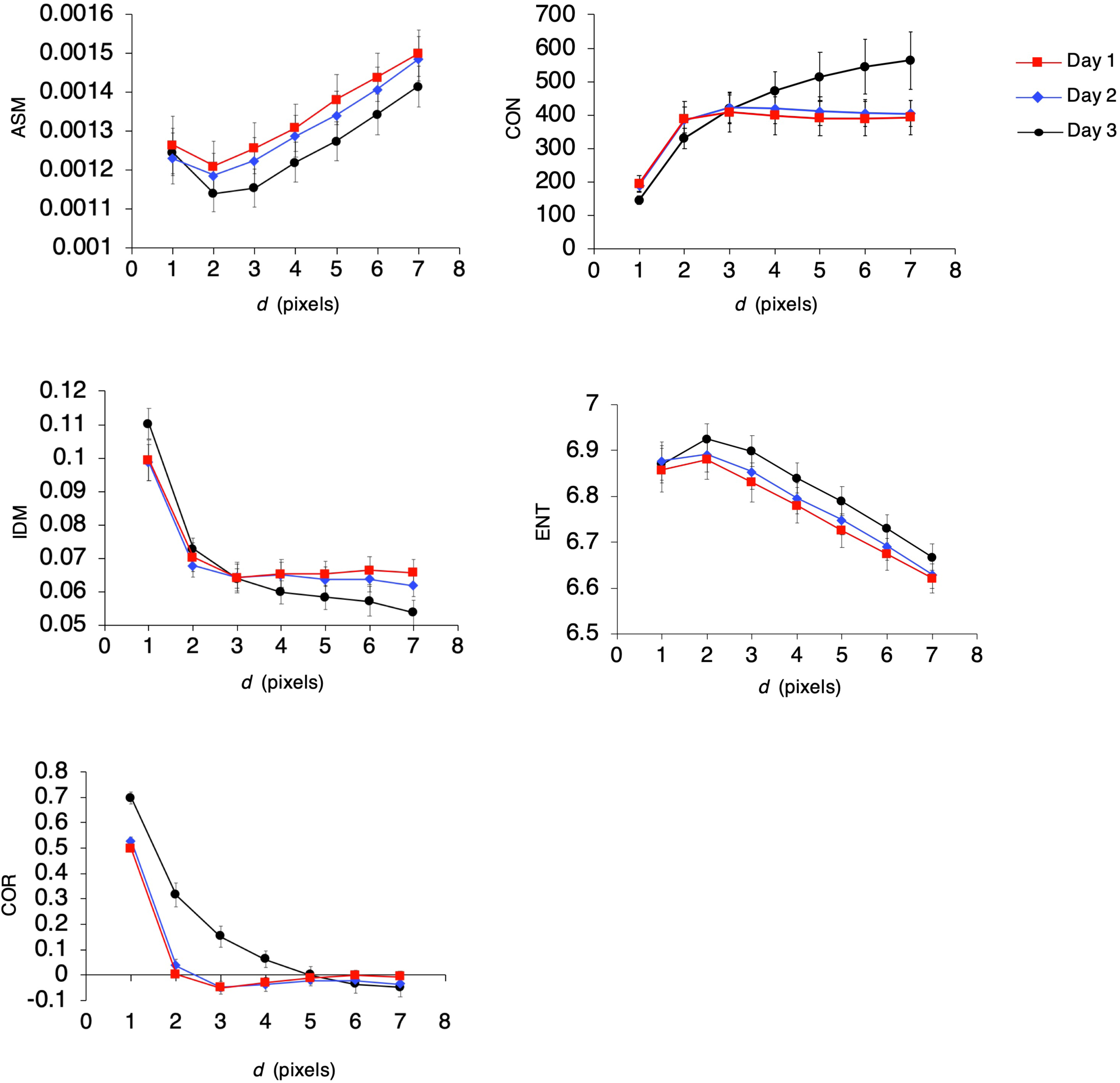
Comparison of texture features based on Gray-Level Co-Occurrence Matrix (GLCM) between Day 1, Day 2, and Day 3 oocytes. Key texture features based on GLCM of Day 1, Day 2, and Day 3 were calculated (n = 12 animals each age group, pooled from two experiments). The curve of the mean texture feature as a function of distance *d*; *θ* = 135 degrees. Error bars indicate SEM. ASM: *Angular Second Moment*, CON: *Contrast*, IDM: *Inverse Difference Moment*, ENT: *Entropy*, COR: *Correlation*.

**Additional file 4: Figure S4.**
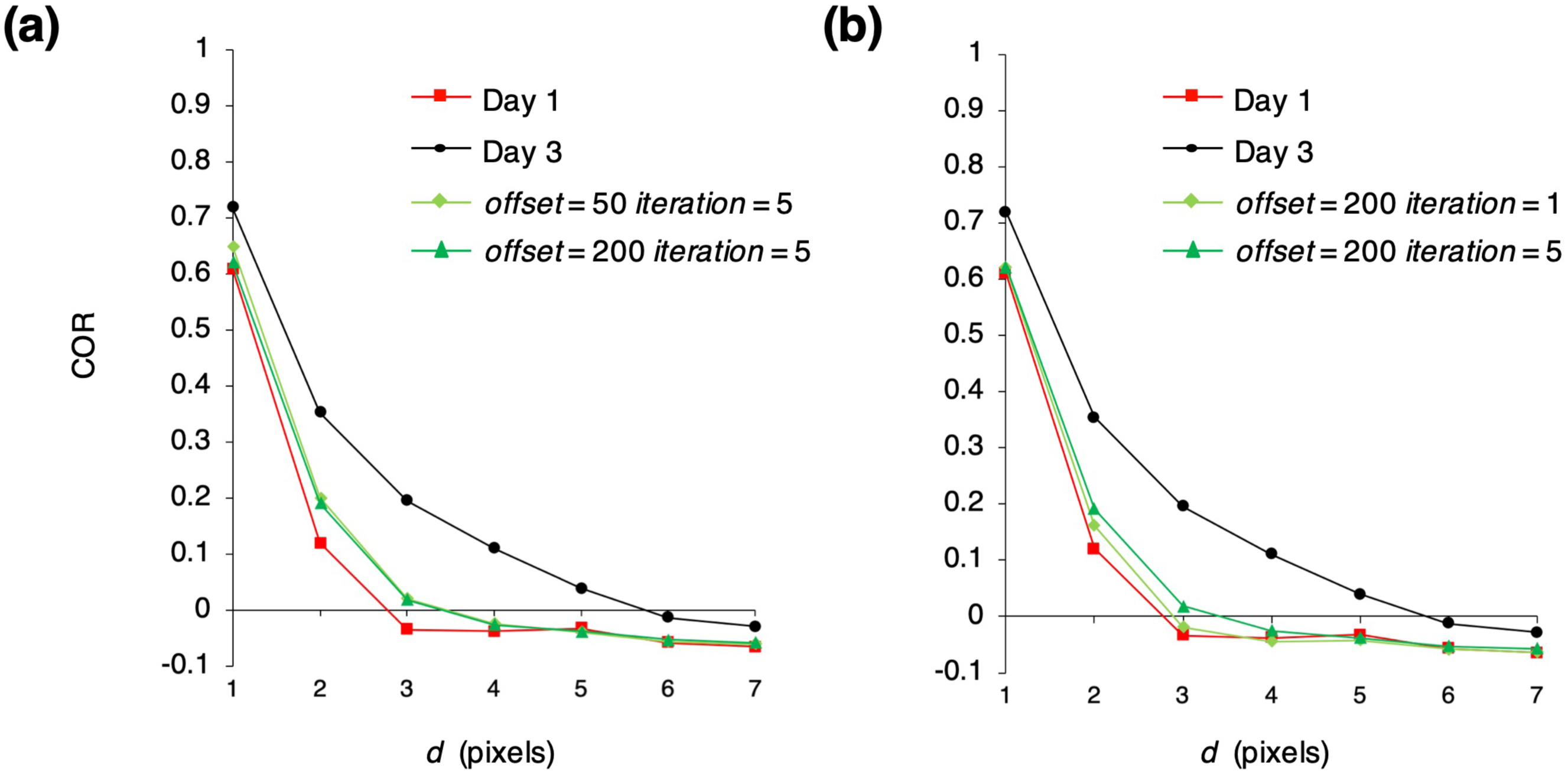
The *Correlation* curve (COR) of cytoplasmic texture calculated for Day 1, Day 3, and Day 3-fied oocytes for the Combination pattern. The mean COR as a function of distance *d* at *θ* = 135 degrees. The COR of the cytoplasmic texture in actual Day 1, actual Day 3, and Day 3-fied oocyte images (n = 10 animals each age group; orientation of the worms is 0 degrees). The parameters for the Combination pattern (*offset* and *iteration*) were set to (a) (50, 5) and (200, 5) or (b) (200, 1) and (200, 5), respectively.

**Additional file 5: Figure S5.**
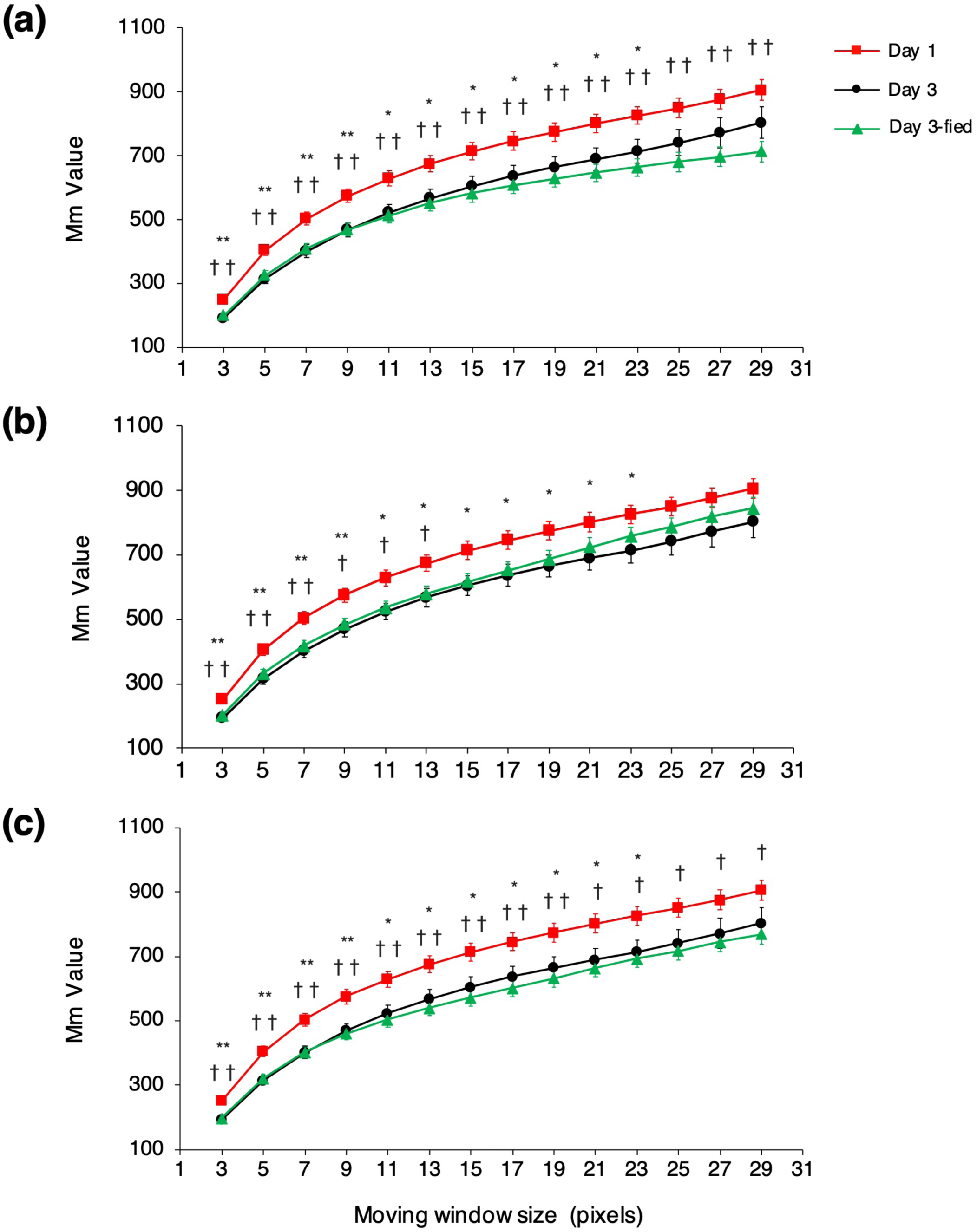
Comparison of Mm Value between actual Day 1, actual Day 3 and Day 3-fied oocyte images after changing the window size. (a-c) Comparison of Mm Value between actual Day 1, actual Day 3 and Day 3-fied oocyte images (n = 10 animals each age group; orientation of the worms is 0 degrees) after changing the window size from 3×3 to 29×29. (a) The parameter for the Smoothed pattern (*offset*) was set to 100. (b) The parameter for the Large Structure pattern (*iteration*) was set to 3. (c) The parameter for the Combination pattern (*offset* and *iteration*) were set to (50, 1). Error bars indicate SEM. Symbols indicate statistical significance (Tukey–Kramer test) between Day 1 and Day 3 (\**P* < 0.05, \*\**P* < 0.01) or Day 1 and Day 3-fied (†*P* < 0.05, ††*P* < 0.01).

## Abbreviations

GLCM: Gray level co-occurrence matrix
COR: *Correlation*
Mm Value: Max-min Value
DIC microscopy: Differential interference contrast microscopy
*C. elegans*: *Caenorhabditis elegans*

## Acknowledgements

We thank Prof. Zhiwei Luo for discussion and general support. The Bristol N2 strain was provided by the *Caenorhabditis elegans* Genetics Center, which is funded by NIH Office of Research Infrastructure Programs (P40 0D010440).

## Availability of data and materials

The oocyte images will be available at the Systems Science of Biological Dynamics database (SSBD) [40], http://ssbd.qbic.riken.jp/set/20200701/

## Authors’ contributions

M.I, J.T and S.O designed research; M.I performed computational experiments; M.I, J.T and H.O performed biological experiments; and M.I. and S.O. wrote the paper.

## Consent for publication

Not applicable.

## Ethics approval and consent to participate

Not applicable.

## References

1. Goud P, Goud A, Van Oostveldt P, Elst Van der Elst J, Dhont M. Fertilization abnormalities and pronucleus size asynchrony after intracytoplasmic sperm injection are related to oocyte postmaturity. Fertil. Steril. 1999; 72: 245–252.

2. Te Velde ER, Pearson PL. The variability of female reproductive ageing. Human Reproduction Update. 2002; 8: 141–154.

3. Miao YL, Kikuchi K, Sun QY, Schatten H. Oocyte aging: cellular and molecular changes, developmental potential and reversal possibility. Human Reproduction Update. 2009; 15: 573–585.

4. Kenyon C. The plasticity of aging: Insights from long-lived mutants. Cell. 2005; 120: 449–460.

5. Andux S, Ellis RE. Apoptosis maintains oocyte quality in aging *Caenorhabditis elegans* females. PLoS Genet. 2008; 4.

6. Klass MR. A method for the isolation of longevity mutants in the nematode *Caenorhabditis elegans* and initial results. Mech. Ageing Dev. 1983; 22: 279–286.

7. Friedman DB, Johnson ET. A mutation in the age-1 gene in *Caenorhabditis elegans* lengthens life and reduces hermaphrodite fertility. Genetics. 1988; 118: 75–86.

8. Kenyon C, Chang J, Gensch E, Rudner A, Tabtiang R. A *C.elegans* mutant that lives twice as long as wild type. Nature. 1993; 366: 461–464.

9. Luo S, Kleemann GA, Ashraf JM, Shaw WM, Murphy CT. TGF-β and insulin signaling regulate reproductive aging via oocyte and germline quality maintenance. Cell. 2010; 143: 299–312.

10. Ortiz De Solorzano C, Costes S, Callahan DE, Parvin B, Barcellos-Hoff MH. Applications of quantitative digital image analysis to breast cancer research. Microsc. Res. Tech. 2002; 59: 119–127.

11. Glotsos D, Spyridonos P, Cavouras D, Ravazoula P, Arapantoni Dadioti P, Nikiforidis G. An image-analysis system based on support vector machines for automatic grade diagnosis of brain-tumour astrocytomas in clinical routine. Med. Inform. Internet Med. 2005; 30: 179–193.

12. Losa GA, Castelli C. Nuclear patterns of human breast cancer cells during apoptosis: characterisation by fractal dimension and co-occurrence matrix statistics. Cell Tissue Res. 2005; 322: 257–267.

13. Masseroli M, Bollea A, Forloni G. Quantitative morphology and shape classification of neurons by computerized image analysis. Comput. Methods Programs Biomed. 1993; 41: 89–99.

14. Chen S, Zhao M, Wu G, Yao C, Zhang J. Recent advances in morphological cell image analysis. Comput. Math. Methods Med. 2012; e101536.

15. Smitha P, Shaji L, Mini MG. A review of medical image classification techniques. Int. Conf. VLSI, Commun. Intrumrnataiom. 2011; 34–38.

16. Angermueller C, Pärnamaa T, Parts L, Stegle O. Deep learning for computational biology. Mol. Syst. Biol. 2016; 12: 878.

17. Haralick RM, Shanmugam K, Dinstein IH. Textural features for image classification. IEEE Trans. Syst. Man. Cybern. 1973; 6: 610–621.

18. Sabino DMU, da Fontoura Costa L, Rizzatti EG, Zago MA. A texture approach to leukocyte recognition. Real-Time Imaging. 2004; 10, 205–216.

19. Castellano G, Bonilha L, Li LM, Cendes F. Texture analysis of medical images. Clin. Radiol. 2004; 59: 1061–1069.

20. Zulpe N, Pawar V. GLCM textural features for brain tumor classification. Int. J. Comput. Sci. Issues. 2012; 9: 354–359.

21. Sulston JE, Horvitz HR. Post-embryonic cell lineage of the nematode, Caenorhabditis elegans. Dev. Biol. 1977; 56: 110–156.

22. Sulston JE, Schierenberg E, White JG, Thomson JN. The embryonic cell lineage of the nematode *Caenorhabditis elegans*. Dev. Biol. 1983; 100: 64–119.

23. Preza C, Snyder DL, Conchello JA. Theoretical development and experimental evaluation of imaging models for differential-interference-contrast microscopy. J. Opt. Soc. Am. A. 1999; 16: 2185–2199.

24. Cheeks RJ, Canman JC, Gabriel WN, Meyer N, Strome S, Goldstein B. *C. elegans* PAR proteins function by mobilizing and stabilizing asymmetrically localized protein complexes. Curr. Biol. 2004; 14: 851–862.

25. Verbrugghe KJ, Chan RC. Imaging *C. elegans* embryos using an epifluorescent microscope and open source software. J. Vis. Exp. 2011; 49: e2625.

26. Sharrock WJ, Sutherlin ME, Leske K, Cheng TK, Kim TY. Two distinct yolk lipoprotein complexes from *Caenorhabditis elegans*. J. Biol. Chem. 1990; 265: 14422–14431.

27. Kimble J, Sharrock WJ. Tissue-specific synthesis of yolk proteins in *Caenorhabditis elegans*. Dev. Biol. 1983; 96: 189–196.

28. Grant B, Hirsh D. Receptor-mediated endocytosis in the *Caenorhabditis elegans* oocyte. Mol. Biol. Cell. 1999; 10: 4311–4326.

29. Garigan D, Hsu AL, Fraser AG, Kamath RS, Ahringer J, Kenyon C. Genetic analysis of tissue aging in *Caenorhabditis elegans*: a role for heat-shock factor and bacterial proliferation. Genetics. 2002; 161: 1101–1112.

30. Herndon LA, Schmeissner PJ, Dudaronek JM, Brown PA, Listner KM, Sakano Y, et al. Stochastic and genetic factors influence tissue-specific decline in ageing *C. elegans*. Nature. 2002; 419: 808–814.

31. Perez MF, Francesconi M, Hidalgo-Carcedo C, Lehner B. Maternal age generates phenotypic variation in *C. elegans*. Nature. 2017; 552: 106–109.

32. Gems D, de la Guardia Y. Alternative perspectives on aging in *Caenorhabditis elegans*: Reactive oxygen species or hyperfunction? Antioxidants Redox Signal. 2013; 19: 321–329.

33. Murphy CT, McCarroll SA, Bargmann CI, Fraser A, Kamath RS, Ahringer J, Li H, Kenyon C. Genes that act downstream of DAF-16 to influence the lifespan of *Caenorhabditis elegans*. Nature. 2003; 424: 277–283.

34. Seah NE, et al. Autophagy-mediated longevity is modulated by lipoprotein biogenesis. Autophagy. 2016; 12: 261–272.

35. Sönnichsen B, et al. Full-genome RNAi profiling of early embryogenesis in *Caenorhabditis elegans*. Nature. 2005; 434: 462–469.

36. Kyoda K. et al. WDDD: worm developmental dynamics database. Nucleic Acids Res. 2013; 41: D732–D737.

37. Brenner S. The genetics of *Caenorhabditis elegans*. Genetics. 1974; 77: 71–94.

38. Kim E, Sun L, Gabel CV, Fang-Yen C. Long-term imaging of *Caenorhabditis elegans* using nanoparticle-mediated immobilization. PLoS One. 2013; 8: e53419.

39. Samuel AD, Murthy VN, Hengartner MO. Calcium dynamics during fertilization in *C. elegans*. BMC Dev. Biol. 2001; 1: 8.

40. Tohsato Y, Ho KH, Kyoda K, Onami S. SSBD: a database of quantitative data of spatiotemporal dynamics of biological phenomena. Bioinformatics. 2016; 32: 3471–3479.

